## Supplementary Information for "Ribosomal DNA copy number is associated with body mass in humans and other mammals"

**Supplementary Material**

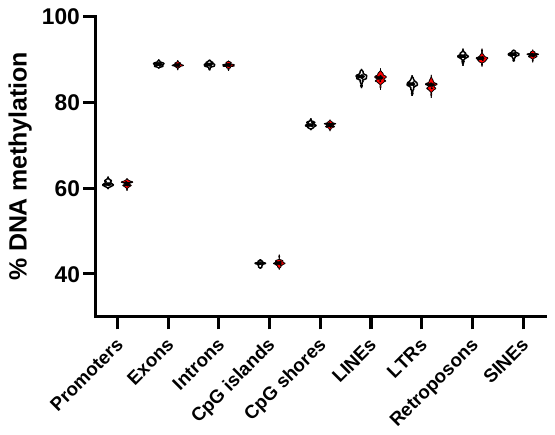

**Figure S1. Methylation of classes of genomic features in lean and obese groups of the discovery cohort.** Average methylation was calculated for all CpG sites captured within reads mapping to the above specific annotations. No features were considered to be differentially methylated between lean and obese groups with Multiple Mann-Whitney tests (P_adj_ <0.01). The lean group is represented by clear and obese by red violin plots.

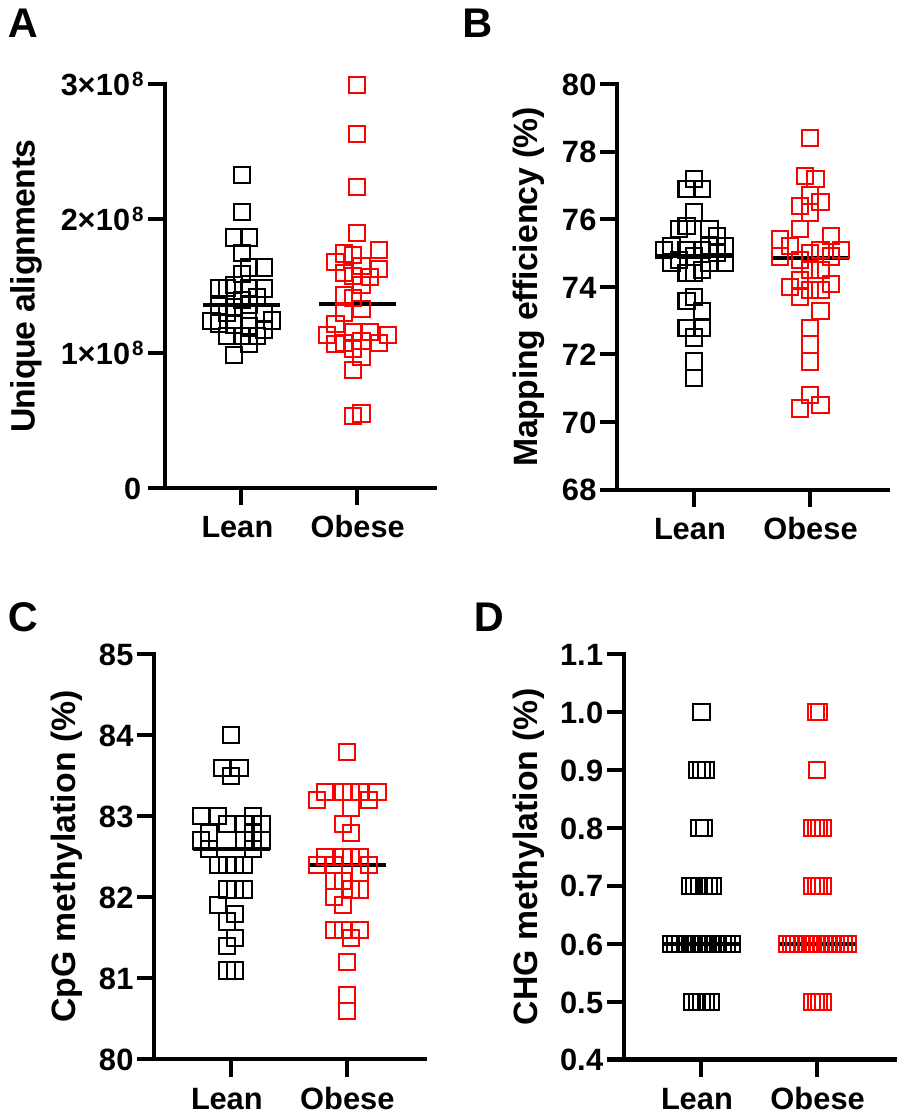

**Figure S2. Pairwise comparison of sequencing statistics from the age-matched, multi-ethnic discovery cohort after exclusions based on data quality control. A)** Reads uniquely mapped to the whole genome + rDNA consensus are not different between lean or obese males (p = 0.6671, Mann-Whitney test). **B)** Mapping efficiency is not variable between lean or obese males (p = 0.8031, Mann-Whitney test). **C)** Global CpG methylation estimates are not different between lean or obese males (p = 0.6103, Mann-Whitney test). **D)** Non-CpG (CHG) methylation is not different between lean or obese males (p = 0.9956, Mann-Whitney test). Throughout lean (BMI<25 kg/m^2^, n=31, black) or obese (BMI>30 kg/m^2^, n=32, red).

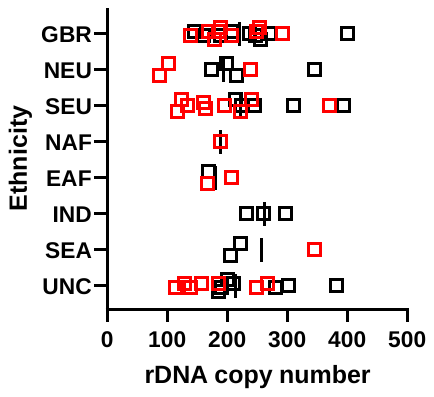

**Figure S3. rDNA copy number broken down by ethnicity in the age-matched mixed ethnicity discovery cohort.** Throughout lean (BMI<25 kg/m^2^, n=31, black) or obese (BMI>30 kg/m^2^, n=32, red). GBR (White British), NEU (Northern European), SEU (Southern European), NAF (North African), EAF (East African), IND (Indian), SEA (South East Asian), UNC (unclassified). The trend for lower rDNA copy number reaches significance in SEU (p=0.0420, Mann-Whitney test) and UNC (p=0.0379, Mann-Whitney test).

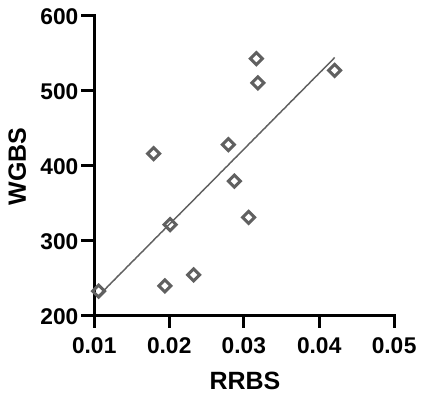

**Figure S4. Cross-validation of copy-number estimation methods.** WGBS data and RRBS data generated from human LCLs analysed with the respective copy number estimation approaches for each data type show a positive correlation (Spearman r = 0.7727, p = 0.0074, n = 11).

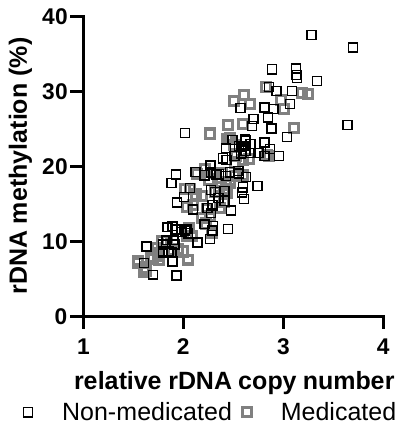
**Figure S5. rDNA copy number and methylation are highly correlated in the adipose tissue from the validation (METSIM) cohort.** All samples (Spearman r=0.8830, p<0.0001, n=169). Non-medicated only (Spearman r=0.8803, p<0.0001, n=100). Medicated only (Spearman r=0.9059, p<0.0001, n=69).

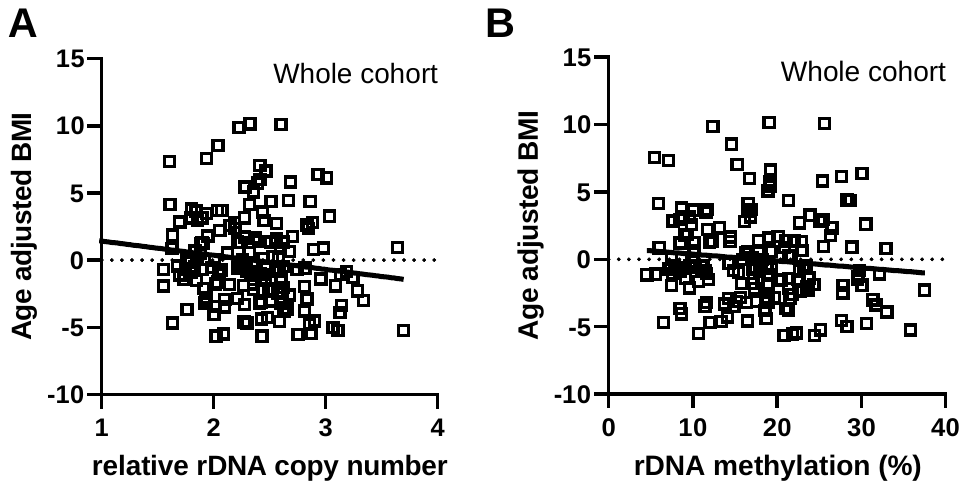
**Figure S6. rDNA copy number negatively correlates with age-adjusted BMI but methylation does not. (A)** rDNA copy number (Spearman r=-0.1657, p=0.0313, n=169) **(B)** rDNA methylation (Spearman r=-0.1238, p=0.1087, n=169).

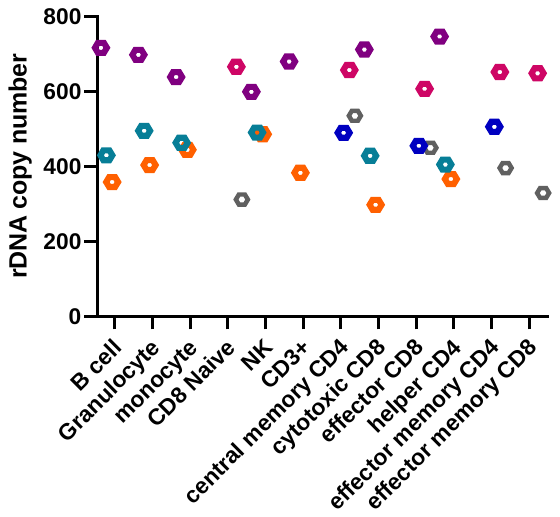
**Figure S7. rDNA copy number does not vary among different blood cell types.** rDNA copy number was estimated using published whole genome sequencing data^1^. Individual donors with multiple purified blood cell types available are represented by different colours.

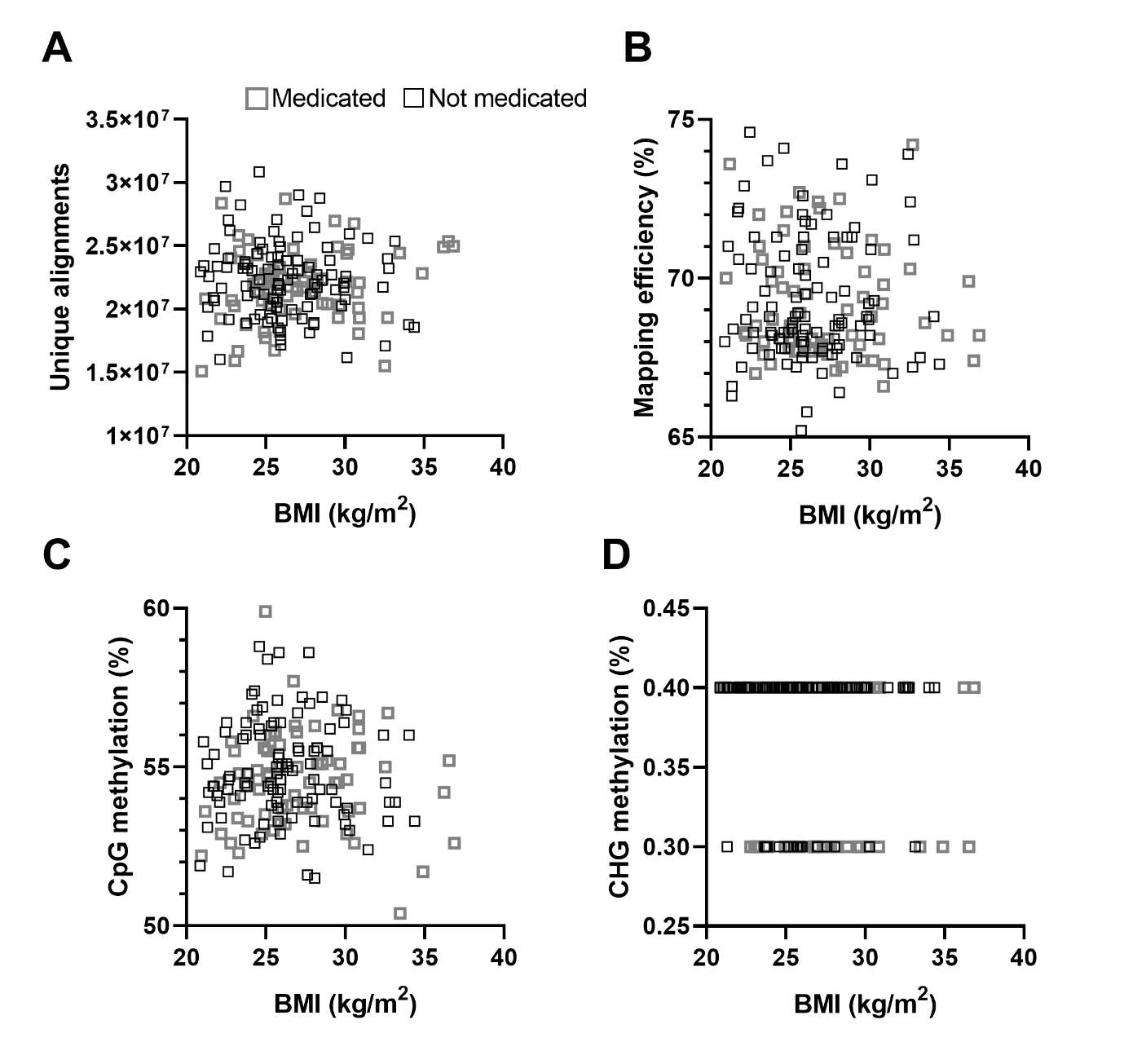

**Figure S8. BMI-association for sequencing statistics for the adipose tissue from the METSIM cohort after exclusions based on data quality control.** Medicated individuals are represented in grey, individuals not taking any medication are represented in black. **A)** Reads uniquely mapped to the whole genome + rDNA consensus are not correlated with BMI (Medicated: Spearman r = 0.1583, p = 0.1940, n = 69 not medicated: Spearman r = -0.06787, p = 0.5022, n = 100). **B)** Mapping efficiency is not correlated with BMI (Medicated: Spearman r = -0.08227, p = 0.5016, n= 69, not medicated: Spearman r = -0.01277, p = 0.8997, n = 100). **C)** Global CpG methylation estimates are not correlated with BMI (Medicated: Spearman r = 0.09548, p = 0.4351, n= 69, not medicated: Spearman r = -0.02857, p = 0.7778, n = 100). **D)** Non-CpG (CHG) methylation is not correlated with BMI (Medicated: Spearman r = -0.1065, p = 0.3837, n= 69, not medicated: Spearman r = 0.02075, p = 0.8376, n = 100).

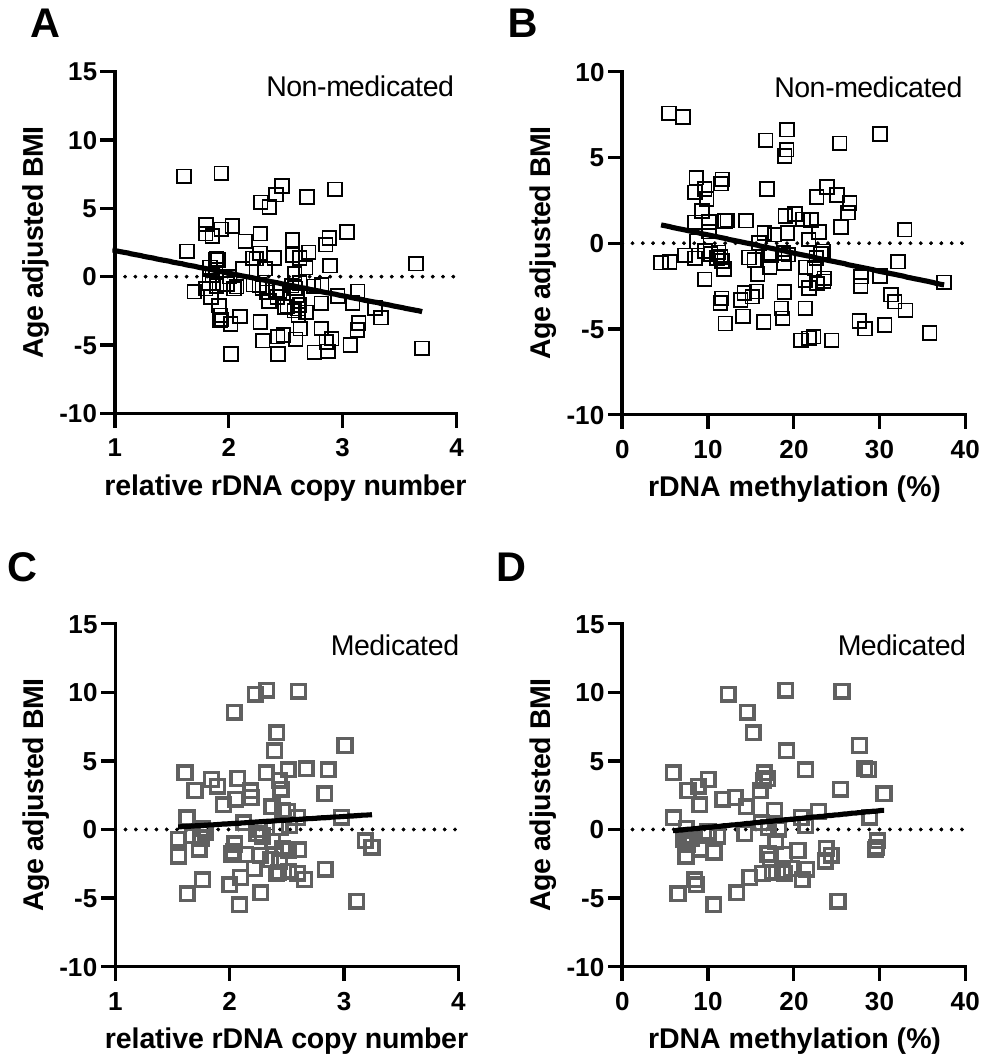
**Figure S9. rDNA copy number and methylation correlate with age-adjusted BMI only in the non-medicated group. (A)** Relative rDNA copy number is negatively correlated with age-adjusted BMI in the non-medicated group (Spearman r=-0.2865, p=0.0039, n=100). **(B)** rDNA methylation is negatively correlated with age-adjusted BMI in the non-medicated group (Spearman r=-0.2487, p=0.0126, n=100). **(C)** Relative rDNA copy number is not correlated with age-adjusted BMI in the non-medicated group (Spearman r=0.0619, p=0.6132, n=69). **(D)** rDNA methylation is not correlated with age-adjusted BMI in the non-medicated group (Spearman r=-0.0832, p=0.4968, n=69)

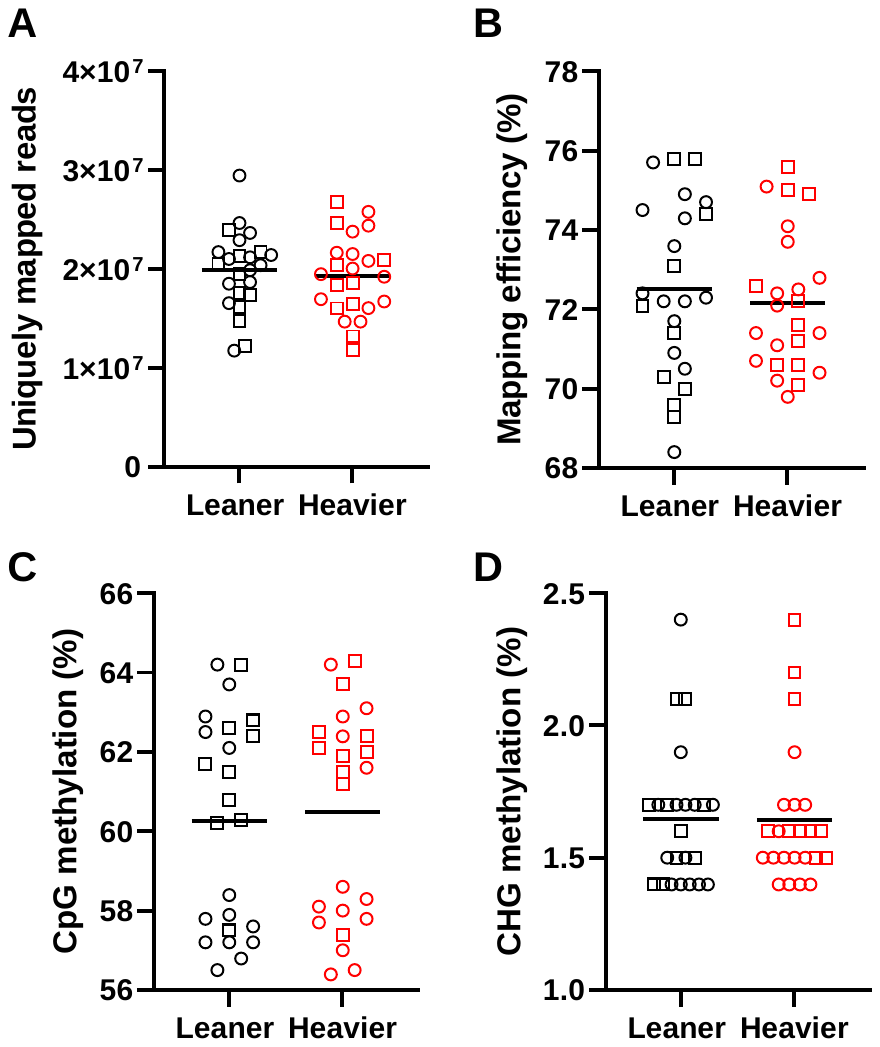
**Figure S10. Pairwise comparison of sequencing statistics for single ethnicity, monozygotic twins discordant for BMI after exclusions based on data quality control. A)** Reads uniquely mapped to the whole genome + rDNA consensus are not different between leaner or heavier twins (p=0.2182, n=24, Wilcoxon matched-pairs signed rank test, Pairing: Spearman r=0.4843, p=0.0082). **B)** Mapping efficiency is not variable between leaner or heavier twins (p=0.2101, n=24, Wilcoxon matched-pairs signed rank test, Pairing: Spearman r=0.5727, p=0.0017). **C)** Global CpG methylation estimates are not different between leaner or heavier twins (p=0.2311, n=24, Wilcoxon matched-pairs signed rank test, Pairing: Spearman r=0.8520, p<0.0001). **D)** Non-CpG (CHG) methylation is not different between leaner or heavier twins (p=0.9414, n=24, Wilcoxon matched-pairs signed rank test, Pairing: Spearman r=0.5859, p=0.0013). Sex of twins is indicated (Male=square, Female=circle).

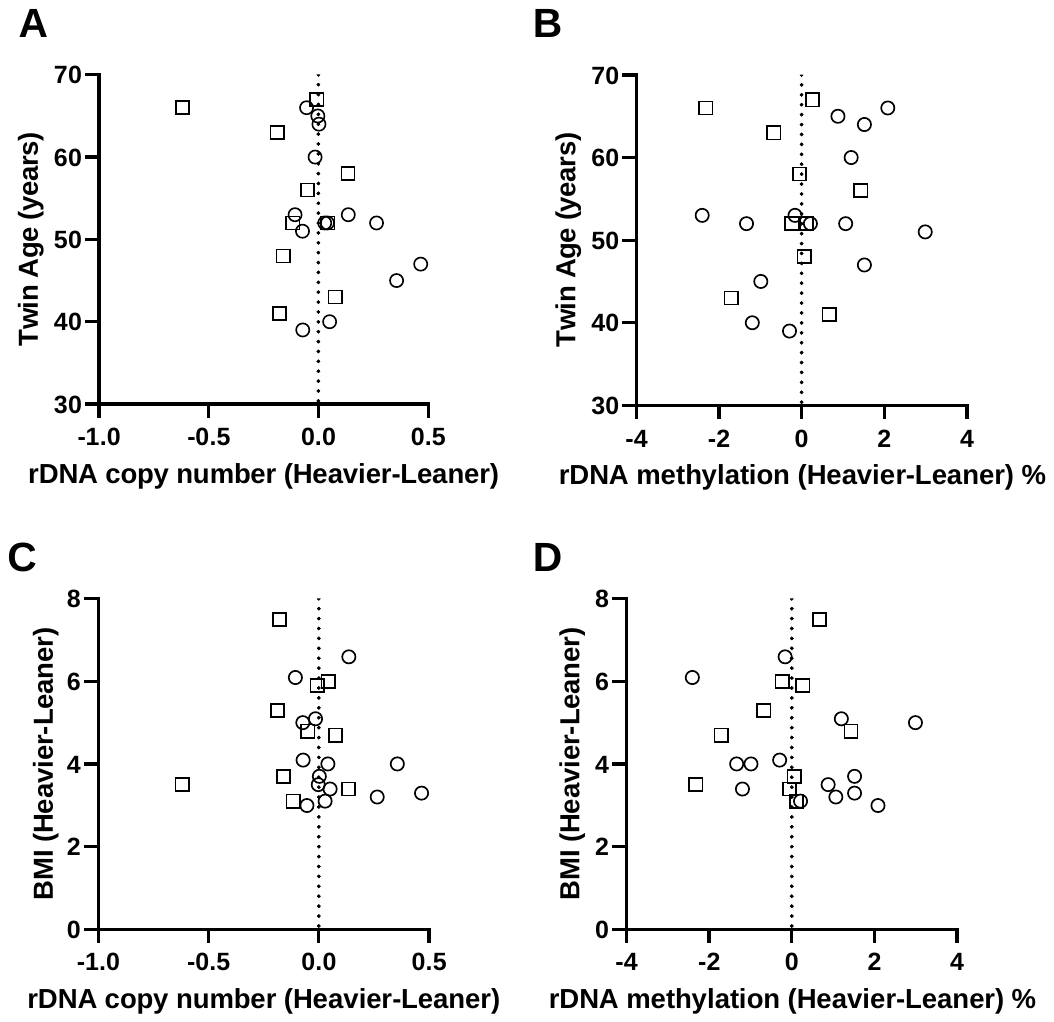
**Figure S11. Age and BMI discordance are not associated with the magnitude of between-twin differences in rDNA copy number or methylation.** Female twin pairs are indicated by circles, male twin pairs by squares. **(A)** There is no correlation between the age of twin pairs and the between-twin difference in estimated relative copy number (Spearman r = -0.2267, p = 0.2867, n = 24). **(B)** There is no correlation between the age of twin pairs and the between-twin difference in rDNA methylation (Spearman r = 0.2141, p = 0.3152, n = 24). **(C)** There is no correlation between the BMI discordance of twin pairs and the between-twin difference in estimated relative copy number (Spearman r = -0.2128, p = 0.3180, n = 23). **(B)** There is no correlation between the BMI discordance of twin pairs and the between-twin difference in rDNA methylation (Spearman r = -0.2224, p = 0.2962, n = 24).

**Figure S12. rDNA copy number and methylation are highly correlated in the blood from the single ethnicity monozygotic twin cohort.** Males are indicated by squares, females by circles (Spearman r=0.8564, p<0.0001, n=48).
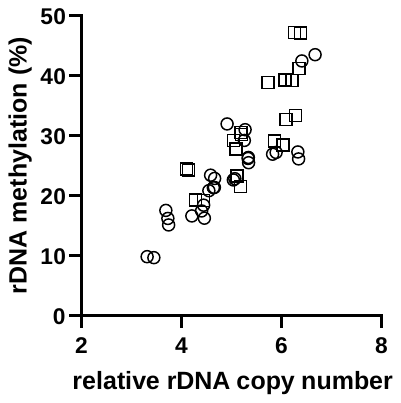

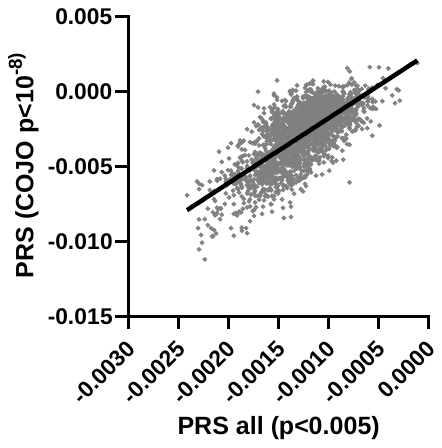
**Figure S13. Correlation of the polygenic risk scores for BMI in the 1000 Genomes Project samples calculated using SNVs that produce the best model for explaining rDNA variance (no COJO, P<0.005) or the COJO filtered SNVs previously used^2^.** Spearman r=0.6734, p<0.0001, n=2390.

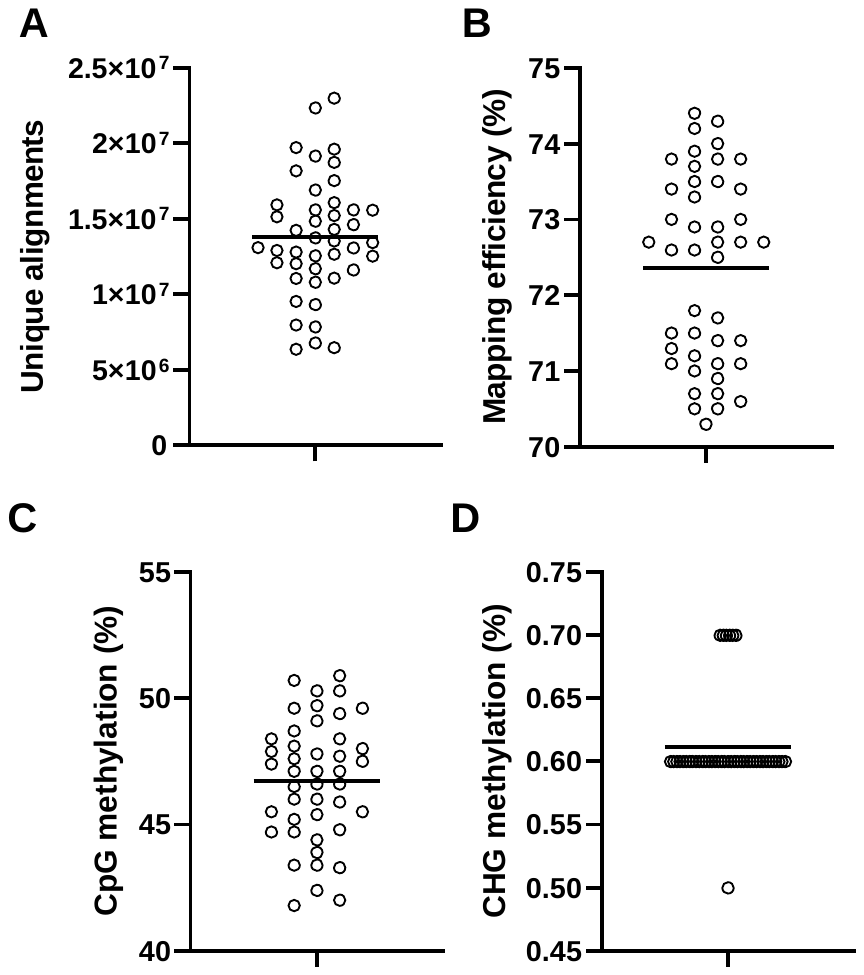
**Figure S14. Distribution of sequencing statistics for RRBS data from liver of Sprague-Dawley rats after exclusions based on data quality control.**

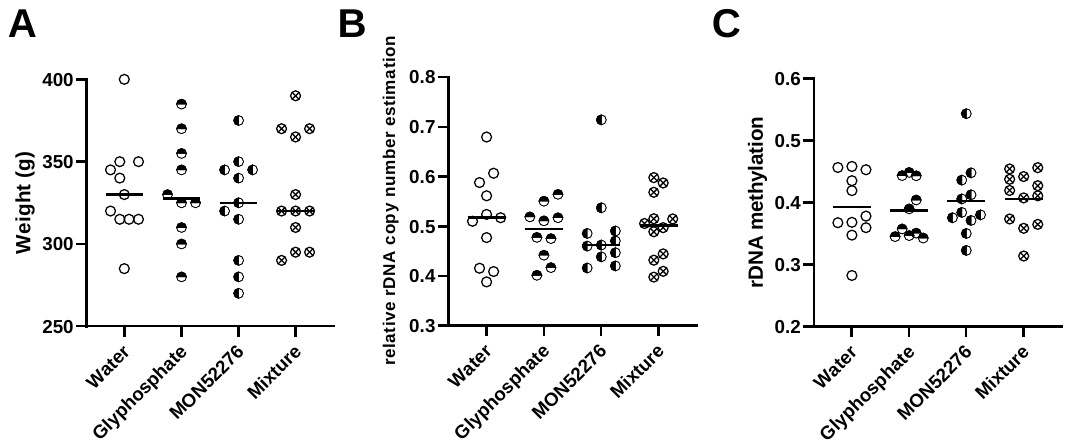

**Figure S15. Chemical treatments administered to female Sprague Dawley rats in the original study do not associate with changes in body mass, rDNA copy number or methylation. (A)** The body mass at the end of the treatment period is not altered by any treatment compared to water (Kruskal-Wallis test, p=0.9352). **(B)** The rDNA copy number is not changes in any treatment compared to water (Kruskal-Wallis test, p=0.7205). **(C)** The rDNA methylation is not changes in any treatment compared to water (Kruskal-Wallis test, p=0.7231).

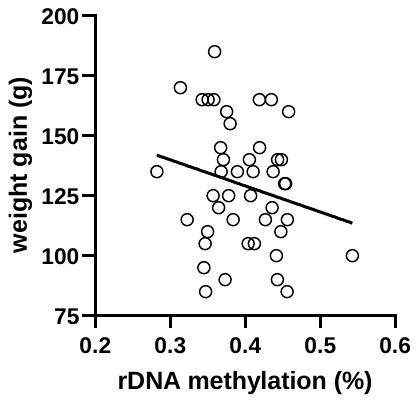
**Figure S16. The weight gained throughout the study period is not significantly correlated with methylation at rDNA.** Spearman r=-0.1598, p=0.3002, n=44.

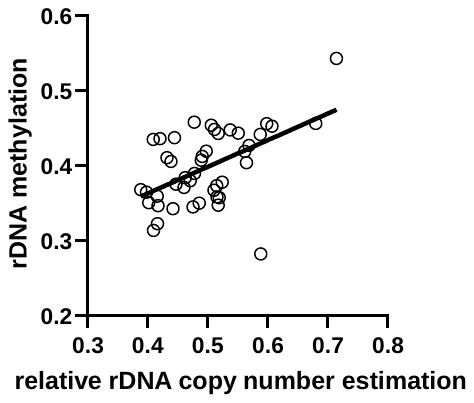
**Figure S17. rDNA copy number and methylation are correlated in the liver of Sprague Dawley rats.** Spearman r=0.4620, p=0.0016, n=44.

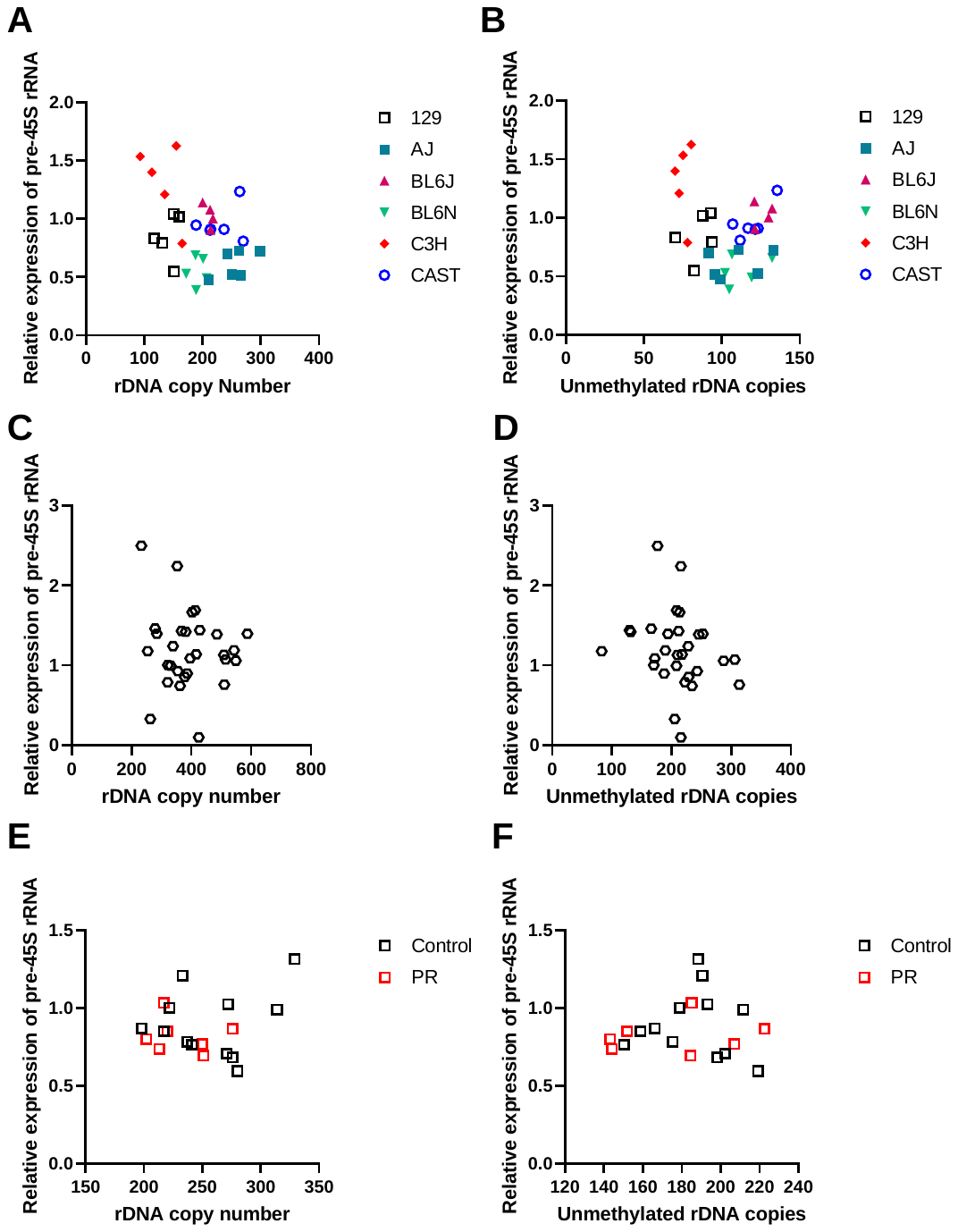
**Fig S18. Nascent 45S-rRNA transcript abundance does not correlate with total rDNA copy number or the number of unmethylated copies present in adult tissues.** Kidney tissue across different mouse strains **(A)** Spearman r=-0.3074, p=0.0926, n=31 **(B)** Spearman r=-0.1266, p=0.4973, n=31. Human LCLCs **(C)** Spearman r=-0.04680, p=0.8095, n=29, **(D)** Spearman r=-0.3340, p=0.0766, n=29. From liver of C57BL/6J mice **(E)** Spearman r=-0.08827, p=0.7193, n=19 **(F)** Spearman r=-0.0009, p=0.9975, n=19.

**Table S1. Summary metabolic data for the lean and obese male cohort.** Indicators of Type 2 Diabetes are from collected data and defined according to the recommendations of the World Health Organisation of HbA1c(%)> 6.5 for Diabetic and 6<HbA1c(%)<6.5 for pre-diabetic ^3^. S.D= standard deviation.

| **Trait** | **Lean** | **Obese** | **Count** | **Mann-Whitney test** |
| --- | --- | --- | --- | --- |
|  | Mean ± S.D. | Mean ± S.D | (lean, obese) | P value |
| BMI (kg/m^2^) | 22.37 ± 1.73 | 33.77 ± 3.06 | 30,31 | P < 0.0001 |
| Age (years) | 35.66 ± 5.27 | 37.38 ± 5.01 | 30,31 | P = 0.1151 |
| Waist circumference (cm) | 81.85 ± 6.66 | 111.7 ± 9.36 | 30,31 | P < 0.0001 |
| Systolic BP (mmHg) | 118.5 ± 10.90 | 133.2 ± 11.28 | 30,31 | P < 0.0001 |
| Diastolic BP (mm/Hg) | 75.74 ± 8.46 | 83.55 ± 8.72 | 30,31 | P = 0.0003 |
| C-reactive protein (mg/L) | 1.28 ± 1.94 | 2.40 ± 2.69 | 30,31 | P = 0.0004 |
| Fasting Glucose (mmol/L) | 4.66 ± 1.18 | 4.53 ± 1.27 | 30,31 | P = 0.5258 |
| HbA1c (%) | 5.03 ± 0.98 | 5.38 ± 0.29 | 30,31 | P = 0.0205 |
| Fasting insulin (mIU/L) | 4.86 ± 3.39 | 12.20 ± 7.97 | 30,30 | P < 0.0001 |
| HOMA-IR | 1.02 ± 0.77 | 2.75 ± 1.95 | 30,30 | P < 0.0001 |
| Total cholesterol (mmol/L) | 4.77 ± 0.97 | 5.15 ± 0.95 | 30,31 | P = 0.1039 |
| Triglycerides (mmol/L) | 1.09 ± 0.59 | 1.69 ± 0.93 | 30,30 | P = 0.0029 |
| HDL (mmol/L) | 1.47 ± 0.29 | 1.29 ± 0.24 | 30,31 | P = 0.0010 |
| LDL (mmol/L) | 2.85 ± 0.90 | 3.17 ± 0.77 | 30,31 | P = 0.1246 |
| Cholesterol:HDL | 3.37 ± 0.94 | 4.10 ± 1.23 | 30,31 | P = 0.0022 |
| Self-identified ever smoker | 8 | 6 | 30, 30 |  |
| Indicators of Type 2 Diabetes | 0 | 0 | 30,30 |  |

**Table S2. Average coverage of genomic features included in methylation analyses.** Values presented are average reads/base in each feature group. rDNA values represent the entire rDNA unit, inclusive of the intergenic spacer.

|  | Lean | | Obese | |  |
| --- | --- | --- | --- | --- | --- |
|  | Mean | S.D. | Mean | S.D. | P value |
| rDNA | 109.57 | 39.89 | 100.91 | 80.20 | 0.0622 |
| Promoters | 302.61 | 74.60 | 313.08 | 154.88 | 0.5075 |
| Exons | 41.48 | 10.00 | 42.49 | 18.37 | 0.5524 |
| Introns | 471.24 | 123.11 | 478.60 | 191.47 | 0.7796 |
| CpG Islands | 229.45 | 56.41 | 238.66 | 124.43 | 0.4564 |
| CpG shores | 200.75 | 47.71 | 208.90 | 100.54 | 0.6181 |
| Simple Repeats | 225.58 | 68.67 | 219.46 | 62.77 | 0.6972 |
| tRNA | 31.11 | 7.78 | 32.07 | 14.00 | 0.6014 |
| LINE | 29.55 | 8.40 | 30.04 | 10.94 | 0.9103 |
| LTR | 23.20 | 5.99 | 23.19 | 9.02 | 0.6181 |
| Retroposon | 95.02 | 19.17 | 97.69 | 43.66 | 0.4816 |
| Satellite | 458.42 | 134.56 | 431.55 | 140.06 | 0.4319 |
| SINE | 34.11 | 8.90 | 35.05 | 14.51 | 0.7979 |

**Table S3. Whole genome bisulfite sequencing quality control from blood for mixed ethnicity, age-matched lean and obese males.**

| Sample ID | Group | Unique alignments | | Mapping efficiency | | CpG methylation | | CHG methylation | | Reason for exclusion |
| --- | --- | --- | --- | --- | --- | --- | --- | --- | --- | --- |
| D002 | Obese | 121889583 | | 75.50% | | 83.10% | | 0.80% | |  |
| D010 | Obese | 107928672 | | 74.80% | | 80.80% | | 0.60% | |  |
| D014 | Obese | 114151592 | | 75.10% | | 83.20% | | 1.00% | |  |
| D020 | Lean | 163979620 | | 75.50% | | 83.60% | | 0.60% | |  |
| D035 | Lean | 130412189 | | 74.70% | | 81.50% | | 0.60% | |  |
| D051 | Lean | 107418211 | | 72.80% | | 81.10% | | 0.70% | |  |
| D055 | Obese | 133033119 | | 77.30% | | 82.80% | | 0.50% | |  |
| D059 | Obese | 109536212 | | 74.50% | | 82.20% | | 0.60% | |  |
| D072 | Lean | 128759190 | | **64.60%** | | 84.10% | | 0.50% | | low ME |
| D073 | Lean | 148720197 | | 76.90% | | 83.00% | | 0.50% | |  |
| D085 | Obese | 107653252 | | 73.70% | | 82.30% | | 0.60% | |  |
| D086 | Lean | 124749055 | | 75.80% | | 81.10% | | 0.60% | |  |
| D087 | Obese | 189452637 | | 75.40% | | 83.20% | | 0.60% | |  |
| D092 | Lean | 98921437 | | 74.90% | | 82.70% | | 0.90% | |  |
| D094 | Lean | 158943462 | | 72.80% | | 82.10% | | 0.90% | |  |
| D095 | Lean | 142062334 | | 77.20% | | 83.60% | | 0.60% | |  |
| D115 | Obese | 160149436 | | 76.40% | | 81.20% | | 0.60% | |  |
| D127 | Lean | 139542680 | | 74.80% | | 82.40% | | 0.60% | |  |
| D128 | Obese | 97714662 | | 74.90% | | 82.50% | | 0.70% | |  |
| D134 | Obese | 165223507 | | 77.20% | | 80.60% | | 0.50% | |  |
| D138 | Obese | 176898387 | | 73.90% | | 82.40% | | 0.60% | |  |
| D145 | Lean | 117580854 | | 74.40% | | 81.70% | | 0.60% | |  |
| D155 | Lean | 148819534 | | 75.70% | | 82.90% | | 0.50% | |  |
| D162 | Lean | 113384625 | | 76.90% | | 82.90% | | 0.70% | |  |
| D166 | Lean | 205493367 | | 73.70% | | 84.00% | | 0.60% | |  |
| D178 | Obese | 174273544 | | 74.50% | | 81.60% | | 0.60% | |  |
| D194 | Obese | 103374637 | | 75.00% | | 82.50% | | 0.60% | |  |
| D198 | Lean | 7873261 | | 74.70% | | 82.40% | | 0.60% | | low SR |
| D199 | Obese | 299075667 | | 70.80% | | 81.50% | | 0.80% | |  |
| D205 | Lean | 136046733 | | 75.00% | | 81.80% | | 0.60% | |  |
| D218 | Obese | 141050678 | | 74.90% | | 83.30% | | 0.60% | |  |
| D224 | Lean | 124152683 | | 75.10% | | 82.70% | | 0.90% | |  |
| D233 | Lean | 186239492 | | 73.60% | | 82.40% | | 0.60% | |  |
| D237 | Lean | 174520020 | | 73.30% | | 82.10% | | 0.60% | |  |
| D244 | Lean | 186022986 | | 75.10% | | 82.90% | | 0.50% | |  |
| D255 | Obese | 19753245 | | 74.30% | | 82.10% | | 0.70% | | low SR |
| D257 | Lean | 163799709 | | 74.70% | | 82.60% | | 0.70% | |  |
| D267 | Lean | 148498650 | | 71.80% | | 82.80% | | 0.60% | |  |
| D274 | Lean | 148551360 | | 74.70% | | 83.00% | | 0.60% | |  |
| D277 | Obese | 156781374 | | 76.50% | | 81.90% | | 0.60% | |  |
| D284 | Obese | 223429145 | | 74.00% | | 82.40% | | 0.60% | |  |
| D292 | Obese | 173219549 | | 70.50% | | 83.30% | | 0.70% | |  |
| D300 | Lean | 135458475 | | 75.10% | | 82.40% | | 0.50% | |  |
| D311 | Obese | 162915645 | | 75.20% | | 82.10% | | 0.60% | |  |
| D316 | Obese | 157318163 | | 78.40% | | 82.10% | | 0.50% | |  |
| D318 | Obese | 262818139 | | 76.70% | | 82.00% | | 0.50% | |  |
| D323 | Lean | 130106545 | | 75.70% | | 82.10% | | 0.50% | |  |
| D324 | Obese | 115801708 | | 73.30% | | 83.80% | | 0.80% | |  |
| D330 | Obese | 88042083 | | 76.20% | | 83.30% | | 0.60% | |  |
| D364 | Obese | 168143562 | | 75.70% | | 81.60% | | 0.60% | |  |
| D369 | Lean | 134735086 | | 76.20% | | 82.70% | | 0.70% | |  |
| D373 | Lean | 122313249 | | 75.20% | | 83.50% | | 1.00% | |  |
| D380 | Lean | 112800154 | | 74.40% | | 82.40% | | 0.60% | |  |
| D381 | Lean | 150926816 | | 74.50% | | 81.40% | | 0.80% | |  |
| D383 | Obese | 143521919 | | 74.10% | | 83.30% | | 0.60% | |  |
| D393 | Obese | **21165587** | | 72.70% | | 83.00% | | 1.00% | | low UA |
| D394 | Obese | 55540544 | | 73.90% | | 82.90% | | 0.60% | |  |
| D398 | Obese | 107207176 | | 70.40% | | 82.40% | | 0.80% | |  |
| D401 | Obese | 53706409 | | 71.80% | | 83.30% | | 0.70% | |  |
| D408 | Lean | 121590427 | | 75.20% | | 81.90% | | 0.80% | |  |
| D416 | Obese | 113861012 | | 75.10% | | 81.60% | | 1.00% | |  |
| D419 | Lean | 120448053 | | 75.20% | | 82.60% | | 0.70% | |  |
| D421 | Obese | 116142496 | | 74.20% | | 82.20% | | 0.90% | |  |
| D427 | Lean | 112762449 | | 72.50% | | 83.00% | | 0.70% | |  |
| D453 | Obese | 150967661 | | 72.30% | | 82.50% | | 0.60% | |  |
| D454 | Lean | 232589678 | | 71.30% | | 82.80% | | 0.60% | |  |
| D463 | Obese | 137341511 | | 74.50% | | 85.50% | | **11.30%** | | poor BC |
| D465 | Lean | 193424213 | | 73.30% | | 82.40% | | **1.50%** | | poor BC |
| D520 | Obese | 130173227 | | 72.80% | | 82.50% | | 0.70% | |  |

ME=mapping efficiency, UA= unique alignments, BC= bisulfite conversion efficiency

**Table S4. Ethnic breakdown of lean and obese groups within mixed-ethnicity cohort**

| **Ethnicity** | **Lean** | **Obese** |
| --- | --- | --- |
| White British (GBR) | 9 | 9 |
| Northern European (NEU) | 4 | 3 |
| Southern European (SEU) | 5 | 9 |
| North African (NAF) | 0 | 1 |
| East African (EAF) | 1 | 2 |
| Indian (IND) | 3 | 0 |
| South East Asian (SEA) | 2 | 1 |
| Unclassified (UNC) | 7 | 7 |
| Total | 31 | 32 |

**Table S5. rDNA copy number does not systematically vary by cell type within an individual.** Two-way ANOVA was performed for the effect of cell-type and donor source were performed on the rDNA copy number calculated from published whole genome sequencing data of purified cell populations^1^. Only donor source was shown to be significant.

| **Tissue of origin** | **Cell type (effect on rDNA CN)** | **Donor identity (effect on rDNA CN)** |
| --- | --- | --- |
| All | P = 0.8604 | P = 3.8 x 10^-11^ |
| Blood | P = 0.8414 | P = 7.5 x 10^-8^ |
| Breast | P = 0.7074 | P = 0.0729 |
| Kidney | P = 0.6890 | P = 0.0221 |
| Neuronal | P = 0.1551 | P = 0.0614 |
| Pancreatic | P = 0.8750 | P = 0.0062 |

**Table S6. Summary phenotype data for included individuals from the METSIM cohort.** Summary statistics for individuals classified by whether they take regular medications or not. Indicators of Type 2 Diabetes are from collected data or previous clinical diagnosis and defined according to the recommendations of the World Health Organisation ^3^. S.D= standard deviation. Comparisons that pass the nominal threshold of P < 0.05 are indicated in bold.

| **Trait** | **Medicated**  **n=69** | **Not medicated**  **n=100** | **Mann Whitney test** |
| --- | --- | --- | --- |
|  | Mean ± S.D. | Mean ± S.D. | P value |
| Lean (BMI<25 kg/m^2^) | 31.88% | 35.00% |  |
| Overweight (25<BMI<30 kg/m^2^) | 44.93% | 53.00% |  |
| Obese (BMI>30 kg/m^2^) | 23.19% | 12.00% |  |
| BMI at recruitment (kg/m^2^) | 27.16 ± 3.68 | 26.22 ± 3.11 | P = 0.1488 |
| BMI at biopsy (kg/m^2^) | 27.16 ± 3.76 | 26.14 ± 2.99 | P = 0.2207 |
| Age (years) | 55.03 ± 3.71 | 54.34 ± 4.65 | P = 0.2387 |
| Weight (kg) | 84.26 ± 12.78 | 81.59 ± 10.60 | P = 0.2989 |
| Height (cm) | 176.0 ± 5.43 | 176.2 ± 6.02 | P = 0.6636 |
| Waist circumference (cm) | 98.78 ± 10.58 | 95.00 ± 9.20 | **P = 0.0396** |
| Hip circumference (cm) | 101.1 ± 7.26 | 100.0 ± 5.16 | P = 0.3205 |
| Waist to Hip ratio | 0.98 ± 0.06 | 0.95 ± 0.06 | **P = 0.0046** |
| Systolic BP (mmHg) | 138.9 ± 17.31 | 133.4 ± 12.31 | P = 0.0926 |
| Diastolic BP (mm/Hg) | 90.31 ± 9.93 | 86.97 ± 7.55 | **P = 0.0480** |
| C-reactive protein (mg/L) | 2.55 ± 5.33 | 2.02 ± 2.53 | P = 0.5438 |
| Fasting glucose recruitment (mmol/L) | 5.80 ± 0.50 | 5.76 ± 0.48 | P = 0.7147 |
| Fasting glucose biopsy (mmol/L) | 5.57 ± 0.55 | 5.50 ± 0.44 | P = 0.6698 |
| HbA1c (%) | 5.65 ± 0.33 | 5.62 ± 0.30 | P = 0.4598 |
| Fasting insulin recruitment (mIU/L) | 8.48 ± 4.66 | 6.66 ± 3.57 | **P = 0.0130** |
| HOMA-IR | 2.22 ± 1.27 | 1.72 ± 0.99 | **P = 0.0163** |
| Total cholesterol (mmol/L) | 5.46 ± 0.82 | 5.64 ± 0.87 | P = 0.2102 |
| Triglycerides (mmol/L) | 1.45 ± 0.74 | 1.34 ± 0.95 | P = 0.0527 |
| HDL (mmol/L) | 1.48 ± 0.36 | 1.54 ± 0.43 | P = 0.3180 |
| LDL(mmol/L) | 3.42 ± 0.75 | 3.56 ± 0.72 | P = 0.1873 |
| Cholesterol:HDL | 3.88 ± 0.99 | 3.90 ± 1.05 | P = 0.9358 |
| Self-identified ever smoker | 62.32% | 51.00% |  |
| T2D: Previously diagnosed or HbA1c (%) ≥ 6.5 | 7.25 % | 4.00% |  |
| Prediabetic: 6 ≤ Hb1Ac (%) < 6.5 | 17.38% | 12:00% |  |

**Table S7. Reduced representation bisulfite sequencing quality control for adipose tissue from the METSIM cohort**

| Sample ID | Medicated? | BMI group | Unique alignments | Mapping efficiency | CpG methylation | CHG methylation | Exclusion reason |
| --- | --- | --- | --- | --- | --- | --- | --- |
| SRR4418800 | No | Obese | 11397491 | 44.70% | 56.50% | 0.80% | Low ME, Poor BC |
| SRR4418801 | No | Obese | 12185998 | 45.80% | 59.30% | 0.80% | Low ME, Poor BC |
| SRR4418802 | Yes | Overweight | 12685793 | 45.30% | 58.60% | 0.70% | Low ME, Poor BC |
| SRR4418803 | No | Overweight | 12451091 | 46.20% | 56.20% | 0.70% | Low ME, Poor BC |
| SRR4418804 | Yes | Overweight | 24926233 | 67.40% | 56.80% | 0.40% |  |
| SRR4418805 | No | Overweight | 24121180 | 67.20% | 53.80% | 0.40% |  |
| SRR4418806 | Yes | Overweight | 22116443 | 68.00% | 54.80% | 0.40% |  |
| SRR4418807 | No | Obese | 23992293 | 67.20% | 53.30% | 0.40% |  |
| SRR4418808 | No | Overweight | 22724072 | 68.60% | 55.60% | 0.40% |  |
| SRR4418809 | No | Overweight | 23842362 | 67.70% | 56.70% | 0.40% |  |
| SRR4418810 | No | Lean | 22577129 | 68.40% | 54.20% | 0.40% |  |
| SRR4418811 | No | Overweight | 22711876 | 67.50% | 54.30% | 0.40% |  |
| SRR4418812 | No | Lean | 23248900 | 68.20% | 54.80% | 0.40% |  |
| SRR4418813 | No | Lean | 24388790 | 67.80% | 56.80% | 0.40% |  |
| SRR4418814 | Yes | Lean | 24568038 | 68.00% | 54.80% | 0.40% |  |
| SRR4418815 | No | Lean | 23745582 | 67.60% | 52.70% | 0.40% |  |
| SRR4418816 | No | Lean | 30852946 | 74.10% | 56.20% | 0.40% |  |
| SRR4418817 | Yes | Lean | 37341493 | 73.40% | 57.00% | 0.40% | High UA |
| SRR4418818 | No | Lean | 29706512 | 74.60% | 56.10% | 0.40% |  |
| SRR4418819 | Yes | Obese | 19316243 | 74.20% | 56.70% | 0.40% |  |
| SRR4418820 | No | Obese | 21732241 | 73.90% | 56.00% | 0.40% |  |
| SRR4418821 | No | Lean | 23239957 | 73.70% | 55.90% | 0.40% |  |
| SRR4418822 | No | Overweight | 21716930 | 73.60% | 55.60% | 0.40% |  |
| SRR4418823 | Yes | Obese | 22161137 | 68.80% | 52.90% | 0.30% |  |
| SRR4418824 | No | Overweight | 23517674 | 68.20% | 53.30% | 0.30% |  |
| SRR4418825 | Yes | Overweight | 22469659 | 69.50% | 53.70% | 0.40% |  |
| SRR4418826 | Yes | Overweight | 20345493 | 70.20% | 55.10% | 0.30% |  |
| SRR4418827 | Yes | Lean | 16679134 | 70.60% | 53.40% | 0.30% |  |
| SRR4418828 | Yes | Overweight | 18242632 | 70.30% | 55.70% | 0.30% |  |
| SRR4418829 | No | Overweight | 20544518 | 70.90% | 54.80% | 0.30% |  |
| SRR4418830 | Yes | Lean | 15918411 | 71.00% | 55.50% | 0.30% |  |
| SRR4418831 | Yes | Obese | 20084968 | 69.80% | 56.20% | 0.40% |  |
| SRR4418832 | No | Overweight | 19256709 | 69.40% | 56.30% | 0.40% |  |
| SRR4418833 | No | Lean | 18905051 | 70.20% | 56.40% | 0.40% |  |
| SRR4418834 | Yes | Obese | 19310955 | 70.90% | 55.60% | 0.40% |  |
| SRR4418835 | No | Overweight | 18126124 | 68.10% | 57.00% | 0.40% |  |
| SRR4418836 | Yes | Lean | 18197018 | 67.80% | 55.60% | 0.40% |  |
| SRR4418837 | No | Obese | 18782675 | 68.80% | 56.00% | 0.40% |  |
| SRR4418838 | No | Overweight | 17591557 | 69.50% | 56.40% | 0.40% |  |
| SRR4418839 | No | Overweight | 21506831 | 67.50% | 57.10% | 0.30% |  |
| SRR4418840 | Yes | Overweight | 22326263 | 67.20% | 55.10% | 0.30% |  |
| SRR4418841 | Yes | Overweight | 21718271 | 67.70% | 56.10% | 0.30% |  |
| SRR4418842 | No | Lean | 21063241 | 69.10% | 54.40% | 0.30% |  |
| SRR4418843 | No | Overweight | 11912792 | 43.60% | 54.20% | 1.00% | Low ME, Poor BC |
| SRR4418844 | No | Obese | 13060010 | 44.20% | 55.80% | 1.00% | Low ME, Poor BC |
| SRR4418845 | No | Overweight | 12809373 | 43.50% | 56.00% | 1.00% | Low ME, Poor BC |
| SRR4418846 | No | Lean | 13761080 | 44.30% | 52.80% | 1.00% | Low ME, Poor BC |
| SRR4418847 | No | Overweight | 18924739 | 68.70% | 55.50% | 0.40% |  |
| SRR4418848 | No | Lean | 19580066 | 67.80% | 56.00% | 0.40% |  |
| SRR4418849 | No | Lean | 19186092 | 68.30% | 54.60% | 0.40% |  |
| SRR4418850 | Yes | Overweight | 19327677 | 69.40% | 54.50% | 0.30% |  |
| SRR4418851 | Yes | Lean | 21304708 | 71.50% | 54.30% | 0.40% |  |
| SRR4418852 | No | Overweight | 24895293 | 71.30% | 55.50% | 0.40% |  |
| SRR4418853 | No | Overweight | 23850975 | 71.60% | 56.20% | 0.40% |  |
| SRR4418854 | No | Overweight | 24114842 | 72.00% | 54.20% | 0.40% |  |
| SRR4418855 | Yes | Overweight | 21364622 | 72.50% | 56.30% | 0.40% |  |
| SRR4418856 | Yes | Lean | 20257518 | 72.00% | 54.00% | 0.40% |  |
| SRR4418857 | No | Overweight | 18516318 | 72.60% | 53.70% | 0.40% |  |
| SRR4418858 | Yes | Lean | 20814522 | 73.60% | 53.60% | 0.40% |  |
| SRR4418859 | Yes | Lean | 20679712 | 72.10% | 52.90% | 0.40% |  |
| SRR4418860 | No | Overweight | 18183648 | 71.80% | 54.60% | 0.40% |  |
| SRR4418861 | No | Lean | 26217818 | 71.30% | 54.70% | 0.40% |  |
| SRR4418862 | Yes | Overweight | 24759720 | 72.40% | 57.70% | 0.30% |  |
| SRR4418863 | Yes | Lean | 28364403 | 68.20% | 52.90% | 0.40% |  |
| SRR4418864 | Yes | Overweight | 28717534 | 67.70% | 53.20% | 0.40% |  |
| SRR4418865 | Yes | Obese | 26765972 | 68.10% | 52.60% | 0.40% |  |
| SRR4418866 | No | Lean | 27035190 | 69.10% | 51.70% | 0.40% |  |
| SRR4418867 | No | Lean | 28247676 | 69.60% | 54.10% | 0.40% |  |
| SRR4418868 | No | Overweight | 24926095 | 68.80% | 52.90% | 0.30% |  |
| SRR4418869 | No | Overweight | 27748836 | 69.40% | 53.90% | 0.30% |  |
| SRR4418870 | No | Overweight | 26131180 | 70.30% | 56.40% | 0.30% |  |
| SRR4418871 | Yes | Overweight | 21616514 | 68.40% | 55.50% | 0.40% |  |
| SRR4418872 | Yes | Obese | 19800721 | 70.50% | 15.10% | 0.40% | Low CpGm |
| SRR4418873 | No | Lean | 24016591 | 67.80% | 54.30% | 0.40% |  |
| SRR4418874 | No | Lean | 18995219 | 69.70% | 13.10% | 0.40% | Low CpGm |
| SRR4418875 | No | Overweight | 21173042 | 68.50% | 55.10% | 0.40% |  |
| SRR4418876 | Yes | Overweight | 23007122 | 67.70% | 53.00% | 0.40% |  |
| SRR4418877 | Yes | Overweight | 20414694 | 68.20% | 55.20% | 0.30% |  |
| SRR4418878 | No | Obese | 20750314 | 68.20% | 53.20% | 0.40% |  |
| SRR4418879 | No | Lean | 21179253 | 68.30% | 53.20% | 0.40% |  |
| SRR4418880 | Yes | Obese | 22093712 | 67.30% | 53.70% | 0.40% |  |
| SRR4418881 | No | Lean | 21646540 | 68.70% | 53.40% | 0.40% |  |
| SRR4418882 | No | Overweight | 23042534 | 67.80% | 55.60% | 0.30% |  |
| SRR4418883 | No | Lean | 22343926 | 68.30% | 58.80% | 0.30% |  |
| SRR4418884 | Yes | Lean | 24280483 | 68.10% | 54.90% | 0.40% |  |
| SRR4418885 | No | Overweight | 23163605 | 68.60% | 54.40% | 0.30% |  |
| SRR4418886 | No | Overweight | 15460721 | 72.40% | 15.90% | 0.30% | Low CpGm |
| SRR4418887 | Yes | Overweight | 22922982 | 71.10% | 54.50% | 0.40% |  |
| SRR4418888 | No | Obese | 22600187 | 70.90% | 56.80% | 0.40% |  |
| SRR4418889 | Yes | Obese | 24913137 | 69.90% | 54.20% | 0.40% |  |
| SRR4418890 | No | Obese | 23227914 | 71.20% | 53.90% | 0.40% |  |
| SRR4418891 | No | Overweight | 21525131 | 68.50% | 53.90% | 0.40% |  |
| SRR4418892 | Yes | Lean | 19247768 | 68.30% | 54.50% | 0.40% |  |
| SRR4418893 | No | Overweight | 20188018 | 67.60% | 51.60% | 0.40% |  |
| SRR4418894 | No | Overweight | 19966927 | 68.30% | 53.40% | 0.40% |  |
| SRR4418895 | Yes | Overweight | 16734199 | 72.70% | 56.00% | 0.40% |  |
| SRR4418896 | Yes | Lean | 15089233 | 70.00% | 52.20% | 0.40% |  |
| SRR4418897 | No | Obese | 16177393 | 73.10% | 53.70% | 0.40% |  |
| SRR4418898 | No | Overweight | 13981854 | 71.90% | 48.40% | 0.30% | Low CpGm |
| SRR4418899 | No | Lean | 20683028 | 72.20% | 54.40% | 0.40% |  |
| SRR4418900 | Yes | Overweight | 20475840 | 69.00% | 53.30% | 0.40% |  |
| SRR4418901 | No | Lean | 20907709 | 72.10% | 54.40% | 0.40% |  |
| SRR4418902 | No | Overweight | 18256291 | 74.40% | 33.90% | 0.30% | Low CpGm |
| SRR4418903 | No | Lean | 16025363 | 72.90% | 53.90% | 0.40% |  |
| SRR4418904 | Yes | Obese | 15520718 | 70.30% | 55.00% | 0.40% |  |
| SRR4418905 | No | Obese | 17097725 | 72.40% | 54.50% | 0.40% |  |
| SRR4418906 | Yes | Lean | 12000743 | 75.10% | 32.50% | 0.30% | Low CpGm |
| SRR4418907 | No | Overweight | 20699919 | 62.40% | 54.40% | 0.30% | Low ME |
| SRR4418908 | No | Lean | 19799969 | 61.30% | 53.60% | 0.40% | Low ME |
| SRR4418909 | Yes | Obese | 21718719 | 62.50% | 52.70% | 0.30% | Low ME |
| SRR4418910 | No | Overweight | 18772255 | 63.10% | 50.50% | 0.40% | Low ME |
| SRR4418911 | No | Lean | 22205439 | 63.30% | 52.00% | 0.40% | Low ME |
| SRR4418912 | Yes | Lean | 21114946 | 63.10% | 53.70% | 0.40% | Low ME |
| SRR4418913 | Yes | Lean | 21721579 | 63.50% | 52.40% | 0.40% | Low ME |
| SRR4418914 | No | Overweight | 24720810 | 64.30% | 50.60% | 0.40% | Low ME |
| SRR4418915 | Yes | Obese | 18068891 | 66.60% | 56.60% | 0.30% |  |
| SRR4418916 | No | Overweight | 19085551 | 65.80% | 55.10% | 0.30% |  |
| SRR4418917 | No | Lean | 17871762 | 66.30% | 55.10% | 0.30% |  |
| SRR4418918 | No | Lean | 20164086 | 66.60% | 53.10% | 0.40% |  |
| SRR4418919 | No | Lean | 20119482 | 64.20% | 55.30% | 0.40% | Low ME |
| SRR4418920 | Yes | Overweight | 21713250 | 63.90% | 55.30% | 0.40% | Low ME |
| SRR4418921 | No | Overweight | 20134415 | 65.20% | 55.00% | 0.40% |  |
| SRR4418922 | No | Lean | 14842163 | 68.10% | 13.40% | 0.30% | Low CpGm |
| SRR4418923 | No | Lean | 21322164 | 68.10% | 52.60% | 0.40% |  |
| SRR4418924 | Yes | Obese | 22833253 | 68.20% | 51.70% | 0.30% |  |
| SRR4418925 | Yes | Overweight | 22586230 | 67.70% | 56.00% | 0.30% |  |
| SRR4418926 | No | Overweight | 21991302 | 67.90% | 53.30% | 0.30% |  |
| SRR4418927 | Yes | Lean | 21162434 | 71.90% | 23.90% | 0.30% | Low CpGm |
| SRR4418928 | Yes | Overweight | 21710651 | 67.60% | 52.50% | 0.40% |  |
| SRR4418929 | Yes | Overweight | 21478387 | 67.90% | 55.00% | 0.30% |  |
| SRR4418930 | Yes | Overweight | 22202119 | 68.10% | 53.70% | 0.30% |  |
| SRR4418931 | Yes | Overweight | 22404613 | 70.80% | 55.10% | 0.40% |  |
| SRR4418932 | No | Lean | 23047517 | 71.30% | 57.30% | 0.40% |  |
| SRR4418933 | Yes | Overweight | 23332220 | 71.00% | 55.00% | 0.40% |  |
| SRR4418934 | No | Obese | 23560794 | 72.00% | 56.30% | 0.40% | extreme BMI |
| SRR4418935 | No | Lean | 21820934 | 68.30% | 56.00% | 0.30% |  |
| SRR4418936 | Yes | Lean | 22792840 | 68.10% | 53.50% | 0.30% |  |
| SRR4418937 | No | Lean | 22957284 | 68.00% | 51.90% | 0.40% |  |
| SRR4418938 | Yes | Overweight | 22345042 | 67.80% | 54.10% | 0.30% |  |
| SRR4418939 | Yes | Lean | 24030376 | 67.00% | 52.60% | 0.30% |  |
| SRR4418940 | Yes | Overweight | 23536708 | 67.10% | 53.70% | 0.30% |  |
| SRR4418941 | No | Obese | 25584994 | 67.00% | 52.40% | 0.40% |  |
| SRR4418942 | Yes | Lean | 25817645 | 67.60% | 52.30% | 0.30% |  |
| SRR4418943 | Yes | Overweight | 22467195 | 71.60% | 23.60% | 0.30% | Low CpGm |
| SRR4418944 | Yes | Overweight | 26973992 | 67.90% | 54.50% | 0.40% |  |
| SRR4418945 | No | Obese | 25361985 | 67.50% | 53.90% | 0.30% |  |
| SRR4418946 | No | Overweight | 27067667 | 68.00% | 53.90% | 0.30% |  |
| SRR4418947 | No | Overweight | 21516198 | 68.40% | 54.30% | 0.40% |  |
| SRR4418948 | No | Overweight | 25695633 | 67.00% | 53.90% | 0.30% |  |
| SRR4418949 | No | Obese | 25956842 | 69.30% | 53.00% | 0.30% |  |
| SRR4418950 | Yes | Lean | 24825521 | 68.50% | 48.70% | 0.30% | Low CpGm |
| SRR4418951 | No | Lean | 23527675 | 68.80% | 54.50% | 0.30% |  |
| SRR4418952 | Yes | Obese | 25368291 | 67.40% | 55.20% | 0.30% |  |
| SRR4418953 | Yes | Overweight | 24719201 | 71.70% | 32.70% | 0.30% | Low CpGm |
| SRR4418954 | Yes | Overweight | 27711198 | 69.30% | 49.50% | 0.40% | Low CpGm |
| SRR4418955 | Yes | Obese | 24967414 | 68.20% | 52.60% | 0.40% |  |
| SRR4418956 | No | Overweight | 26439038 | 66.40% | 51.50% | 0.40% |  |
| SRR4418957 | Yes | Lean | 25479679 | 68.70% | 53.30% | 0.30% |  |
| SRR4418958 | Yes | Obese | 24426854 | 68.60% | 50.40% | 0.30% |  |
| SRR4418959 | Yes | Overweight | 19954351 | 67.90% | 53.20% | 0.40% |  |
| SRR4418960 | Yes | Lean | 18758453 | 67.30% | 54.40% | 0.40% |  |
| SRR4418961 | No | Obese | 18581228 | 67.30% | 53.30% | 0.40% |  |
| SRR4418962 | No | Lean | 19220782 | 68.10% | 57.40% | 0.40% |  |
| SRR4418963 | No | Overweight | 20243906 | 68.80% | 57.10% | 0.40% |  |
| SRR4418964 | No | Overweight | 18778947 | 68.50% | 54.60% | 0.40% |  |
| SRR4418965 | No | Overweight | 22044949 | 69.20% | 56.40% | 0.40% |  |
| SRR4418966 | No | Overweight | 17174878 | 70.10% | 54.60% | 0.40% |  |
| SRR4418967 | No | Overweight | 21711748 | 68.70% | 53.50% | 0.40% |  |
| SRR4418968 | No | Overweight | 21879664 | 68.00% | 55.40% | 0.40% |  |
| SRR4418969 | No | Overweight | 21228517 | 67.80% | 54.00% | 0.40% |  |
| SRR4418970 | No | Lean | 21579482 | 69.40% | 44.90% | 0.40% | Low CpGm |
| SRR4418971 | No | Overweight | 20247263 | 68.80% | 54.50% | 0.40% |  |
| SRR4418972 | No | Overweight | 18994895 | 68.40% | 58.40% | 0.40% |  |
| SRR4418973 | Yes | Overweight | 20842264 | 68.90% | 56.10% | 0.40% |  |
| SRR4418974 | Yes | Overweight | 22091967 | 71.80% | 57.70% | 0.60% | Poor BC |
| SRR4418975 | No | Overweight | 29022324 | 70.50% | 55.50% | 0.40% |  |
| SRR4418976 | No | Overweight | 22282595 | 71.30% | 57.20% | 0.40% |  |
| SRR4418977 | No | Lean | 23420332 | 71.00% | 55.80% | 0.40% |  |
| SRR4418978 | No | Overweight | 19234095 | 72.00% | 57.20% | 0.40% |  |
| SRR4418979 | Yes | Obese | 21331776 | 69.20% | 55.60% | 0.40% |  |
| SRR4418980 | Yes | Lean | 20700983 | 68.50% | 55.80% | 0.40% |  |
| SRR4418981 | Yes | Overweight | 20414428 | 69.60% | 54.70% | 0.40% |  |
| SRR4418982 | Yes | Lean | 22201289 | 69.70% | 54.60% | 0.40% |  |
| SRR4418983 | Yes | Overweight | 21027227 | 68.10% | 53.80% | 0.40% |  |
| SRR4418984 | No | Overweight | 22039111 | 67.70% | 55.30% | 0.40% |  |
| SRR4418985 | Yes | Lean | 17749323 | 68.50% | 59.90% | 0.40% |  |
| SRR4418986 | No | Overweight | 20810972 | 68.90% | 54.30% | 0.40% |  |
| SRR4418987 | Yes | Overweight | 40958483 | 67.00% | 55.10% | 0.40% | High UA |
| SRR4418988 | No | Overweight | 45542663 | 66.50% | 54.30% | 0.40% | High UA |
| SRR4418989 | No | Overweight | 22824028 | 69.70% | 54.90% | 0.40% |  |
| SRR4418990 | Yes | Lean | 23655658 | 69.90% | 54.80% | 0.40% |  |
| SRR4418991 | No | Overweight | 28767011 | 69.60% | 54.30% | 0.40% |  |
| SRR4418992 | No | Lean | 25267639 | 70.70% | 52.80% | 0.40% |  |
| SRR4418993 | No | Lean | 24787887 | 70.60% | 55.40% | 0.40% |  |
| SRR4418994 | No | Lean | 23304128 | 70.30% | 56.40% | 0.40% |  |
| SRR4418995 | Yes | Lean | 21979659 | 70.20% | 56.60% | 0.40% |  |
| SRR4418996 | Yes | Obese | 24397582 | 71.20% | 54.60% | 0.40% |  |
| SRR4418997 | No | Overweight | 25344748 | 71.30% | 58.60% | 0.40% |  |
| SRR4418998 | No | Overweight | 23825766 | 71.70% | 55.10% | 0.40% |  |
| SRR4418999 | No | Overweight | 23309361 | 71.30% | 58.60% | 0.40% |  |
| SRR4419000 | Yes | Overweight | 19608976 | 72.20% | 56.30% | 0.40% |  |
| SRR4419001 | No | Lean | 24743412 | 67.30% | 56.90% | 0.40% |  |
| SRR4419002 | No | Overweight | 22309722 | 67.50% | 55.00% | 0.40% |  |
| SRR4419003 | No | Lean | 23375138 | 67.20% | 54.10% | 0.40% |  |
| SRR4419004 | Yes | Obese | 24691974 | 67.40% | 53.60% | 0.40% |  |

ME= mapping efficiency, BC = bisulfite conversion efficiency, UA= unique alignments, CpGm= Global CpG methylation

**Table S8. rDNA copy number and methylation correlation statistics with phenotypes in the mixed ethnicity cohort and the METSIM cohort.** Phenotypic variables available for the mixed-ethnicity obese male cohort and the METSIM (Finnish) male cohort were correlated with rDNA copy number (top panel) and rDNA methylation (lower panel). Correlations with a P value <0.05 are in bold. Only individuals with data for all variables were included.

|  | Mixed ethnicity cohort (n=62) | | METSIM cohort (n=100) | |
| --- | --- | --- | --- | --- |
|  | Correlation with rDNA copy number | | Correlation with rDNA copy number | |
| Trait | Spearman r | P value | Spearman r | P value |
| BMI (kg/m^2^) | **-0.3655** | **0.0035** | **-0.2992** | **0.0025** |
| Age (years) | -0.0705 | 0.5860 | 0.1636 | 0.1039 |
| Waist circumference (cm) | **-0.3607** | **0.0040** | **-0.2487** | **0.0126** |
| Systolic BP (mmHg) | -0.1307 | 0.3113 | **-0.2023** | **0.0436** |
| Diastolic BP (mm/Hg) | -0.1444 | 0.2628 | **-0.2539** | **0.0109** |
| C-reactive protein (mg/L) | **-0.4614** | **0.0002** | -0.1153 | 0.2531 |
| Fasting Glucose (mmol/L) | -0.0936 | 0.4691 | -0.0691 | 0.4948 |
| HbA1c (%) | -0.2285 | 0.0741 | 0.0110 | 0.9135 |
| Fasting insulin (mIU/L) | **-0.3110** | **0.0139** | -0.1652 | 0.1004 |
| HOMA-IR | **-0.3001** | **0.0178** | -0.1684 | 0.0941 |
| Total cholesterol (mmol/L) | -0.0060 | 0.9629 | -0.1459 | 0.1457 |
| Triglycerides (mmol/L) | -0.0305 | 0.8142 | -0.0273 | 0.7878 |
| HDL (mmol/L) | 0.2482 | 0.0517 | -0.1168 | 0.2470 |
| LDL(mmol/L) | -0.0777 | 0.5484 | -0.1214 | 0.2287 |
| Cholesterol:HDL | -0.0758 | 0.5582 | 0.0686 | 0.4979 |
|  | Mixed ethnicity cohort (n=62) | | METSIM cohort (n=100) | |
|  | Correlation with rDNA methylation | | Correlation with rDNA methylation | |
| Trait | Spearman r | P value | Spearman r | P value |
| BMI (kg/m^2^) | -0.2352 | 0.0658 | **-0.2518** | **0.0115** |
| Age (years) | -0.0022 | 0.9862 | 0.0545 | 0.5903 |
| Waist circumference (cm) | -0.2261 | 0.0772 | **-0.2080** | **0.0378** |
| Systolic BP (mmHg) | -0.1689 | 0.1894 | **-0.2245** | **0.0247** |
| Diastolic BP (mm/Hg) | -0.1501 | 0.2442 | -0.1791 | 0.0746 |
| C-reactive protein (mg/L) | **-0.4344** | **0.0004** | -0.1507 | 0.1344 |
| Fasting Glucose (mmol/L) | -0.1880 | 0.1434 | 0.0243 | 0.8101 |
| HbA1c(%) | -0.2077 | 0.1053 | 0.1004 | 0.3203 |
| Fasting insulin (mIU/L) | **-0.2559** | **0.0447** | -0.1223 | 0.2255 |
| HOMA-IR | -0.2460 | 0.0539 | -0.1172 | 0.2453 |
| Total cholesterol (mmol/L) | -0.0074 | 0.9544 | -0.1254 | 0.2137 |
| Triglycerides (mmol/L) | 0.0068 | 0.9582 | -0.0613 | 0.5450 |
| HDL (mmol/L) | 0.1552 | 0.2283 | -0.0639 | 0.5276 |
| LDL(mmol/L) | -0.0721 | 0.5777 | -0.1082 | 0.2838 |
| Cholesterol:HDL | -0.0608 | 0.6389 | 0.0064 | 0.9493 |

**Table S9. Reduced representation bisulfite sequencing quality control from blood for single ethnicity, mixed sex monozygotic twin cohort.**

| **Sample ID** | **Unique Alignments** | **Mapping Efficiency** | **CpG methylation** | **CHG methylation** | **Exclusion Criteria** |
| --- | --- | --- | --- | --- | --- |
| 1009 | 29439353 | 68.4 | 57.9 | 1.4 |  |
| 1010 | 21521971 | 69.8 | 57 | 1.4 |  |
| 1021 | 18530199 | 74.5 | 56.8 | 1.5 |  |
| 1022 | 16715634 | 72.1 | 56.5 | 1.5 |  |
| 2015 | 19477145 | 72.8 | 56.4 | 1.4 |  |
| 2016 | 20395274 | 74.9 | 56.5 | 1.7 |  |
| 2089 | 24633598 | 73.6 | 57.6 | 1.4 |  |
| 2090 | 20813319 | 71.1 | 58.3 | 1.4 |  |
| 2107 | 24394164 | 72.4 | 58 | 1.4 |  |
| 2108 | 21700616 | 74.7 | 57.2 | 1.4 |  |
| 2139 | 16037036 | 74.1 | 58.6 | 1.5 |  |
| 2140 | 21009074 | 72.4 | 57.2 | 1.7 |  |
| 2163 | 25795434 | 70.7 | 57.8 | 1.7 |  |
| 2164 | 21424361 | 72.2 | 57.8 | 1.7 |  |
| 2169 | 23772101 | 70.4 | 58.1 | 1.7 |  |
| 2170 | 23641412 | 72.3 | 57.2 | 1.7 |  |
| 2193 | 23956282 | 71.4 | 57.5 | 1.7 |  |
| 2194 | 24633996 | 72.2 | 57.4 | 1.6 |  |
| 2221 | 19885958 | 70.9 | 64.2 | 1.4 |  |
| 2222 | 14682001 | 70.2 | 63.1 | 1.5 |  |
| 2259 | 21661700 | 72.5 | 57.7 | 1.7 |  |
| 2260 | 22910003 | 71.7 | 58.4 | 1.7 |  |
| 2303 | 19217537 | 75.1 | 64.2 | 1.9 |  |
| 2304 | 21174858 | 75.7 | 62.9 | 1.9 |  |
| 2307 | 20489230 | 75.9 | 63.2 | **4.8** | Poor BC |
| 2308 | 22662553 | 76.3 | 62.7 | 1.8 | matched twin |
| 3003 | 18034310 | 76.3 | 53.1 | 2 | matched twin |
| 3004 | **73003456** | 73.7 | 48.9 | 2 | high UA |
| 3021 | 16148628 | 75.8 | 64.2 | 2.1 |  |
| 3022 | 16013988 | 75.6 | 63.7 | 2.2 |  |
| 3071 | 16810651 | 76.9 | 62.2 | 2.1 | matched twin |
| 3072 | 21772136 | 76.4 | 64 | **4** | Poor BC |
| 3073 | 20934974 | 70.6 | 61.9 | 1.6 |  |
| 3074 | 21359829 | 72.1 | 60.3 | 1.5 |  |
| 3081 | 24801336 | 76.3 | 62.8 | 1.6 | matched twin |
| 3082 | 15661669 | 76 | 61.7 | **4.1** | Poor BC |
| 3097 | 13171611 | 75 | 61.5 | 2.1 |  |
| 3098 | 19491514 | 70 | 60.2 | 1.4 |  |
| 3111 | 20516682 | 69.6 | 61.5 | 1.5 |  |
| 3112 | 18346793 | 71.6 | 62.5 | 1.6 |  |
| 4049 | 16682305 | 75.4 | 60.9 | **4.9** | Poor BC |
| 4050 | 15617841 | 75.8 | 61.4 | 1.9 | matched twin |
| 4055 | 26753987 | 70.1 | 62 | 1.5 |  |
| 4056 | 14786620 | 70.3 | 62.6 | 1.7 |  |
| 4077 | 16960558 | 71.4 | 62.9 | 1.5 |  |
| 4078 | 18674677 | 70.5 | 62.5 | 1.5 |  |
| 4079 | 12244902 | 75.8 | 62.4 | 2.1 |  |
| 4080 | 11865310 | 74.9 | 64.3 | 2.4 |  |
| 4101 | 20416572 | 70.6 | 61.2 | 1.6 |  |
| 4102 | 17368592 | 69.3 | 60.8 | 1.7 |  |
| 4141 | 17569837 | 74.4 | 62.8 | 1.6 |  |
| 4142 | 18597707 | 72.6 | 62.4 | 1.6 |  |
| 4163 | 20065956 | 71.4 | 61.6 | 1.6 |  |
| 4164 | 11741549 | 74.3 | 63.7 | 2.4 |  |
| 5025 | 21711330 | 73.1 | 61.7 | 1.4 |  |
| 5026 | 16477973 | 71.2 | 62.1 | 1.5 |  |
| 5047 | 14678199 | 73.7 | 62.4 | 1.5 |  |
| 5048 | 16555305 | 72.2 | 62.1 | 1.4 |  |
| 5057 | 13920776 | 75.4 | 62.7 | 2.4 | matched twin |
| 5058 | 15060362 | 75.1 | 65.9 | **4.3** | Poor BC |

UA= unique alignments, BC= bisulfite conversion efficiency. The co-twin was excluded if the data from the other did not pass quality control.

**Table S10. Summary data of cohort characteristics for monozygotic twin data.**

|  |  |  | Leaner | Heavier |  |
| --- | --- | --- | --- | --- | --- |
|  | Number of MZ twin pairs | Range of BMI discordance (kg/m^2^) | Average BMI ± S.D (kg/m^2^) | Average BMI ± S.D (kg/m^2^) | Average Age ± S.D (years) |
| Female | 14 | 3-6.6 | 22.84 ± 3.44 | 27.99 ± 4.15 | 52.8 ± 8.6 |
| Male | 10 | 3.1-7.5 | 21.76 ± 2.54 | 26.55 ± 2.43 | 54.6 ± 9.1 |

**Table S11. Reduced representation bisulfite sequencing quality control from liver of Sprague-Dawley rats.**

| Sample ID | Unique Alignments | Mapping efficiency | CpG methylation | CHG methylation | Reason for exclusion |
| --- | --- | --- | --- | --- | --- |
| SRR12598979 | 12097904 | 73.00% | 49.10% | 0.70% |  |
| SRR12598980 | 6357932 | 71.10% | 47.40% | 0.60% |  |
| SRR12598981 | 12809462 | 72.50% | 45.20% | 0.60% |  |
| SRR12598982 | 14389177 | **67.60%** | 46.00% | 0.60% | low ME |
| SRR12598983 | 13077304 | 72.70% | 46.60% | 0.60% |  |
| SRR12598984 | 13742668 | 74.20% | 43.30% | 0.60% |  |
| SRR12598985 | 14234046 | 72.70% | 48.10% | 0.60% |  |
| SRR12598986 | 7841866 | 70.60% | 47.60% | 0.50% |  |
| SRR12598987 | 11059713 | 73.40% | 44.80% | 0.60% |  |
| SRR12598988 | 16072649 | 71.50% | 45.40% | 0.60% |  |
| SRR12598989 | 15131082 | 73.90% | 44.70% | 0.60% |  |
| SRR12598990 | 12546316 | 71.50% | 41.80% | 0.60% |  |
| SRR12598991 | 13514588 | **69.20%** | 48.90% | 0.70% | low ME |
| SRR12598992 | 19154644 | 74.30% | 45.50% | 0.60% |  |
| SRR12598993 | **78493837** | 70.90% | 46.60% | 0.60% | high UA |
| SRR12598994 | 11071671 | 73.80% | 44.70% | 0.60% |  |
| SRR12598995 | 14854162 | 70.50% | 46.60% | 0.60% |  |
| SRR12598996 | 15588742 | 71.80% | 47.10% | 0.60% |  |
| SRR12598997 | 14309296 | 71.40% | 46.00% | 0.60% |  |
| SRR12598998 | 17533671 | 71.40% | 46.00% | 0.60% |  |
| SRR12598999 | 15228607 | 72.90% | 47.70% | 0.70% |  |
| SRR12599000 | 22335933 | 71.30% | 43.90% | 0.60% |  |
| SRR12599001 | 10811083 | 74.40% | 42.00% | 0.60% |  |
| SRR12599002 | 22993349 | 71.20% | 47.10% | 0.60% |  |
| SRR12599003 | 13529153 | 72.70% | 46.50% | 0.60% |  |
| SRR12599004 | 11498510 | 74.10% | **37.60%** | 0.60% | Low CpGm |
| SRR12599005 | 12660411 | 72.90% | 50.90% | 0.70% |  |
| SRR12599006 | 11622513 | 73.70% | 43.40% | 0.60% |  |
| SRR12599007 | 16900498 | 70.50% | 47.90% | 0.70% |  |
| SRR12599008 | 14618140 | 73.00% | 50.70% | 0.70% |  |
| SRR12599009 | 15580905 | 73.50% | 45.90% | 0.60% |  |
| SRR12599010 | 12041598 | 72.70% | 47.80% | 0.60% |  |
| SRR12599011 | 6465253 | 71.10% | 48.70% | 0.60% |  |
| SRR12599012 | 9320092 | 73.30% | 48.40% | 0.60% |  |
| SRR12599013 | 18744579 | 72.60% | 49.60% | 0.60% |  |
| SRR12599014 | 15929680 | 70.70% | 49.40% | 0.60% |  |
| SRR12599015 | 19606975 | 70.30% | 48.00% | 0.70% |  |
| SRR12599016 | 12537964 | 73.40% | 49.60% | 0.60% |  |
| SRR12599017 | 12907900 | 72.60% | 47.10% | 0.60% |  |
| SRR12599018 | 6771926 | 70.90% | 50.30% | 0.60% |  |
| SRR12599019 | 19709504 | 73.80% | 50.30% | 0.60% |  |
| SRR12599020 | 11692693 | 71.00% | 44.40% | 0.60% |  |
| SRR12599021 | 13419085 | 70.70% | 47.50% | 0.60% |  |
| SRR12599022 | 15601244 | 73.50% | 45.50% | 0.60% |  |
| SRR12599023 | 18171792 | 74.00% | 43.40% | 0.60% |  |
| SRR12599024 | 9536599 | 73.80% | 42.40% | 0.60% |  |
| SRR12599025 | 13102550 | 71.70% | 49.70% | 0.60% |  |
| SRR12599026 | 7977364 | 71.10% | 48.40% | 0.60% |  |

UA= unique alignments, BC= bisulfite conversion efficiency, CpGm= Global CpG methylation

**Table S12. Correlation between weight at each time point with rDNA copy number in the liver of female Sprague Dawley rats.** n=44 at each time point. Measurement with P<0.05 are indicated in bold.

| Measurement week | Spearman r | P value |
| --- | --- | --- |
| 8 | -0.0718 | 0.6432 |
| 9 | -0.1579 | 0.3088 |
| 10 | -0.1990 | 0.1953 |
| 11 | -0.2875 | 0.0585 |
| 12 | -0.2119 | 0.1673 |
| 13 | -0.1972 | 0.1995 |
| 14 | -0.2689 | 0.0775 |
| 15 | -0.2776 | 0.0681 |
| 16 | -0.2294 | 0.1341 |
| 17 | -0.2788 | 0.0669 |
| **18** | **-0.3294** | **0.0290** |
| **19** | **-0.3652** | **0.0148** |

**Table S13. Masked regions in the human reference sequence**

| **Assembly** | **Contig** | **Start** | **End** |
| --- | --- | --- | --- |
| Hg38 | Chr 1 | 91387225 | 91387553 |
| Hg38 | Chr 21 | 8202092 | 8260970 |
| Hg38 | Chr 21 | 8385101 | 8472093 |
| Hg38 | Chr 21 | 8986700 | 9899746 |
| Hg38 | ChrUn_GL000220v1 | 0 | 161802 |
| Hg38 | Chr22_KI270733v1_random | 0 | 179772 |
| Rn7 | Chr 3 | 2231807 | 2240483 |
| Rn7 | chrUn_NW_023637849v1 | 61191 | 69866 |
| Rn7 | chrUn_NW_023637831v1 | 81687 | 90364 |
